# 2-Cl-C.OXT-A Stimulates Contraction through the Suppression of Phosphodiesterase Activity in Human Induced Pluripotent Stem Cell-derived Cardiac Organoids

**DOI:** 10.1101/553826

**Authors:** Takahiro Kitsuka, Manabu Itoh, Sojiro Amamoto, Ken-ichi Arai, Junichi Oyama, Koichi Node, Shuji Toda, Shigeki Morita, Takahiro Nishida, Koichi Nakayama

## Abstract

**Background:** 2-Cl-C.OXT-A (COA-Cl) is a novel synthesized adenosine analog that activates S1P1 receptor (S1P1R) and combines with adenosine A1 receptor (A1R) in G proteins and was shown to enhance angiogenesis and improve the brain function in rat stroke models. However, the role of COA-Cl in hearts remains unclear. COA-Cl, which has a similar structure to xanthine derivatives, has the potential to suppress phosphodiesterase (PDE), which is an important factor involved in the beating of heart muscle.

**Methods and results:** Cardiac organoids with fibroblasts, human induced pluripotent stem cell-derived cardiac myocytes (hiPSC-CMs), and hiPSC-derived endothelial cells (hiPSC-ECs) were cultured until they started beating. The beating and contraction of organoids were observed before and after the application of COA-Cl. COA-Cl significantly increased the beating rate and fractional area change in organoids. To elucidate the mechanism underlying these effects of COA-Cl on cardiac myocytes, pure hiPSC-CM spheroids were evaluated in the presence/absence of Suramin (antagonist of A1R). The effects of COA-Cl, SEW2871 (direct stimulator of S1P1R), two positive inotropes (Isoproterenol [ISO] and Forskolin [FSK]), and negative inotrope (Propranolol [PRP]) on spheroids were assessed based on the beating rates and cAMP levels. COA-Cl stimulated the beating rates about 1.5-fold compared with ISO and FSK, while PRP suppressed the beating rate. However, no marked changes were observed with SEW2871. COA-Cl, ISO, and FSK increased the cAMP level. In contrast, the level of cAMP did not change with PRP or SEW2871 treatment. The results were the same in the presence of Suramin as absence. Furthermore, an enzyme analysis showed that COA-Cl suppressed the PDE activity by half.

**Conclusions:** COA-Cl, which has neovascularization effects, suppressed PDE and increased the contraction of cardiac organoids, independent of S1P1R and A1R. These findings suggest that COA-Cl may be useful as an inotropic agent for promoting angiogenesis in the future.

## Introduction

Positive inotropic agents are considered to enhance the hemodynamic profile in terms of elevating the cardiac output, decreasing the cardiac filling pressure, and improving the organ perfusion in patients with heart disease. In clinical trials in which inotropic agents were administered to promote the contraction of the dysfunctional heart, safety was a concern from the viewpoint of the long-term prognosis [1]. However, in the clinical setting, inotropic agents remain a major treatment for patients in the decompensated phase of severe heart failure. Positive inotropic drugs are useful for increasing the cardiac output in order to resolve various problems associated with heart failure, especially for patients with a low systolic blood pressure or low cardiac output [2].

COA-Cl is a novel nucleic acid analogue that has been found to promote angiogenesis. Vascular endothelial growth factor (VEGF), platelet-derived growth factor (PDGF), and hepatocyte growth factor (HGF) activate tube formation in cultured human vascular endothelial cells [3][4][5]. COA-Cl has also been shown to facilitate the functional recovery by enhancing angiogenesis and synaptogenesis *in vivo* in a model of ischemia [6]. Focusing on the chemical structure of COA-Cl, we hypothesized that the structure would be similar to that of xanthine derivatives, which act as nonspecific inhibitors of PDE [7]. We predicted that COA-Cl would act as an inhibitor of PDE and exert a positive inotropic effect on the heart. In order to evaluate the effect *in vitro*, we used cardiac organoids composed of human-derived cells. Cardiac myocytes and fibroblasts determine the structural, mechanical, and electrical characteristics of the myocardium [8]. Studies of cardiac myocytes and fibroblasts in two-dimensional (2D) cell cultures have provided valuable information on their biology and regulation. However, cells grown on a flat hard surface are exposed to a different environment from the tissue [9].

Human induced pluripotent stem cells (hiPSCs) are expected to be a useful source of cells for drug discovery and autogenous transplantation [10] [11] [12]. We have been developing a scaffold-free bio-fabrication technology using spheroids as a unit to construct three-dimensional (3D) structures [13]. Three-dimensional cardiac structures using pure hiPSC-CM spheroids in particular are expected to be able to be transplanted and improve the cardiac function of patients with heart failure in the future [14] [15]. In the present study, we created a 3D heart model *in vitro* to predict and analyze the action of COA-Cl in human heart.

## Materials and Methods

### Cardiac organoids and pure hiPSC-CMs spheroid assays

Human dermal fibroblasts (HDFBs) (Lonza, Basel, Switzerland) cultured in low-glucose Dulbecco’s Modified Eagle Medium (DMEM) (Wako, Japan) containing fetal bovine serum (FBS) (10%) and penicillin/streptomycin (PSA) (Wako, Japan) were used for organoid fabrication (at passage 6-7). hiPSC-derived endothelial cells (hiPSC-derived ECs) (Lonza) cultured in EGM-2 media (Lonza) were used for organoid fabrication (at passage 3-4). hiPSC-derived cardiac myocytes (hiPSC-CMs; iCell Cardiomyocytes, Cellular Dynamics International-CDI, Madison, WI, USA) were thawed according to the manufacturer’s protocol. Three cell lines were plated and monitored for cardiac organoids by transferring 100 µL of hiPSC-CMs (1×10^4^ cells), HDFBs (5×10^3^ cells), and hiPSC-ECs (5×10^3^ cells) onto a 96-well plate with a spindle-shaped bottom (Sumitomo Bakelite, Tokyo, Japan). Cardiac organoids were incubated for 72 h in a CO_2_ incubator at 37 °C and cultured in DMEM+FBS+PSA. We replaced the medium every 48 h for 10 days. To elucidate the mechanism through which COA-Cl exerts its effects on cardiac myocytes, pure hiPSC-CM spheroids were re-suspended in plating medium. The following cells were plated with 100 µL of 2.5×–3×10^4^ cells per well onto a spindle-shaped bottom 96-well plate (Sumitomo Bakelite). The plate was incubated at 37 °C for 72 h in a CO_2_ incubator. We replaced the medium with DMEM+FBS+PSA every 48 h (Fig. 1).

**Figure 1.**
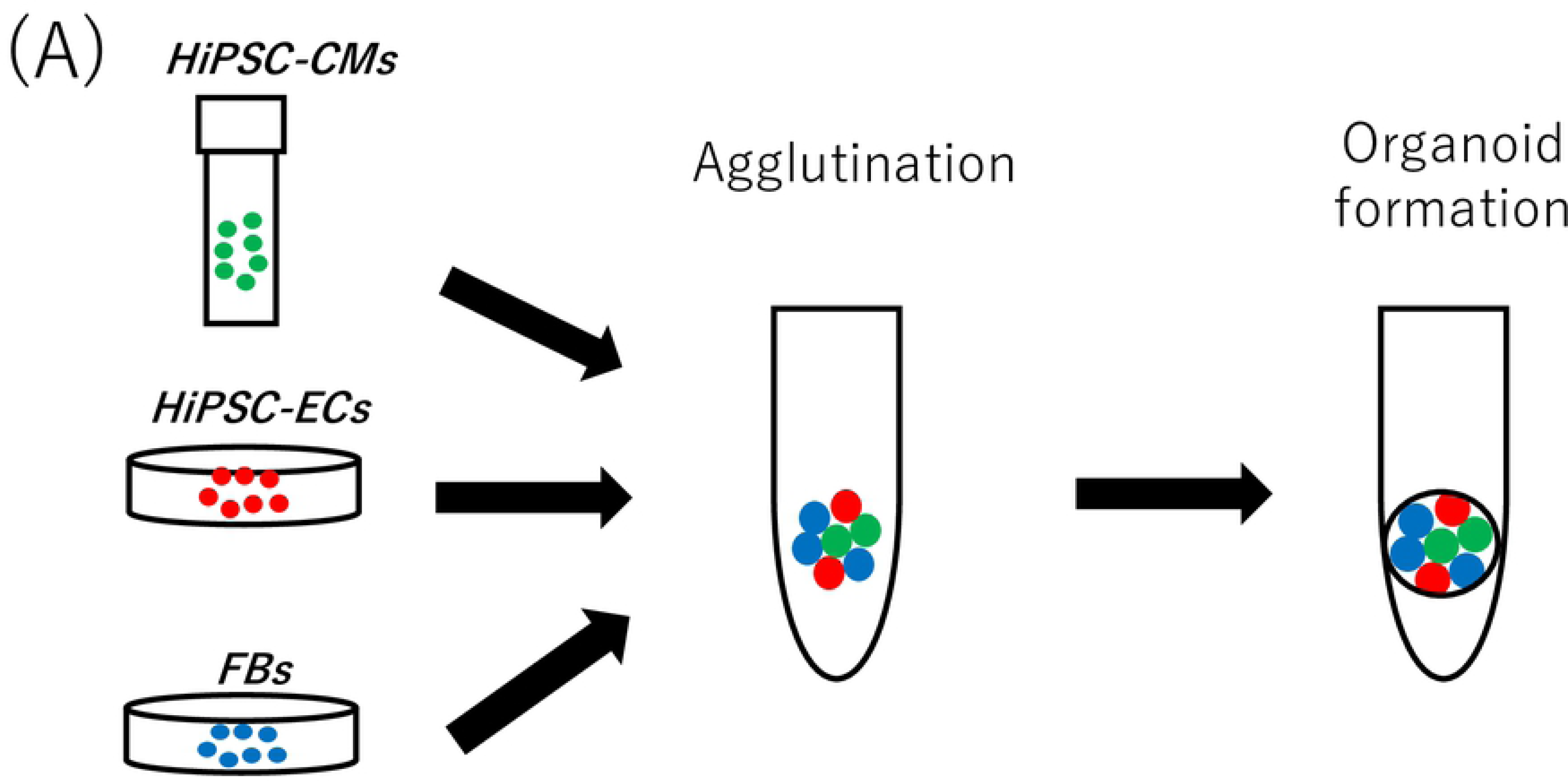
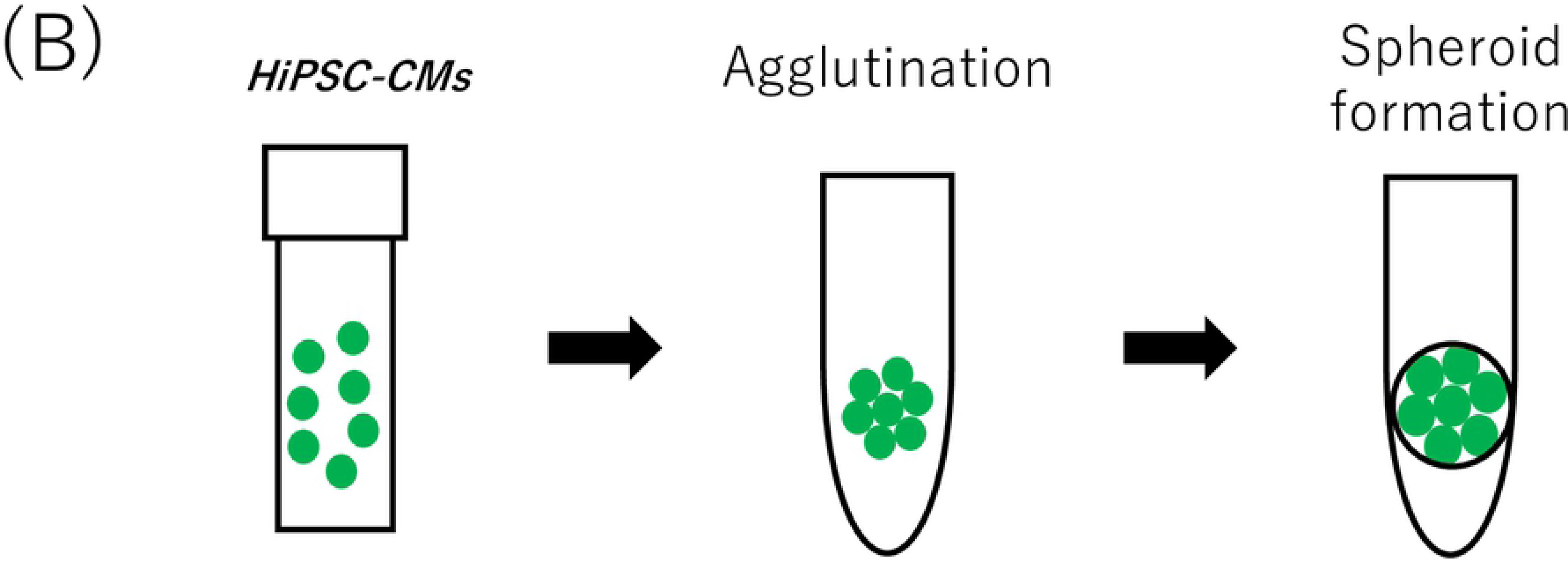
Generation of cardiac organoids in a 96-well spindle plate (A). Generation of pure hiPSC-CM spheroids in a 96-well spindle plate (B).

### The contraction analysis of beating cardiac organoids

We observed beating cardiac organoids before and after the administration of several compounds. Videos of the spontaneously beating cardiac organoids were recorded for 5 min at 37 °C for each condition using a BZX-710 (Keyence, Osaka, Japan). Videos were converted to a series of TIFF format pictures, and the area of the cardiac organoids was measured using the BZ-X Analyzer software program (Keyence). The measured areas were graphed using the Excel software program (Microsoft Excel^®^, Microsoft Co., Ltd., Redmond, WA., USA) to visualize the change in the beating in the fractional area; the beats per minute and contraction amplitude were calculated. The contraction amplitudes were calculated as the fractional area change amplitude between contraction and relaxation.

### The analysis of pure hiPSC-CM spheroids beating

Pure hiPSC-CM spheroids on day 5, after they started beating, were analyzed after replacing the medium with fresh medium and performing incubation at 37 °C for 30 min. We exposed the spheroids to several compounds at different concentrations for 30 min and recorded the findings with a BZX-710. The beating rates were calculated based on moving images. We also detected the onset of the changes in beating after the administration of COA-Cl. We continuously observed the beating spheroids before and after the administration of COA-Cl using fluorescent calcium imaging. The spheroids were incubated for 1 h at 37 °C in loading buffer containing 10 µM fluo4 acetoxymethyl ester (Dojindo, Kumamoto, Japan) and detergents (0.02% Pluronic F-127, and 1.25 mmol/l Probenecid) (Dojindo). The samples were washed with recording buffer (Dojindo) for 20 min at 37 °C, and calcium transients were recorded using fluorescence imaging with an excitation wavelength of 488 nm. The videos were converted to a series of TIFF format pictures, and the relative fluorescence (in relative fluorescence units [RFU]) of the spheroids was measured using the BZ-X Analyzer software program. The measured RFU was graphed to determine the beating profiles of RFU with the Excel software program.

### Intracellular cAMP measurements of pure hiPSC-CM spheroids

First, we measured the half maximal inhibitory concentration (IC50) of COA-Cl with a Lance Ultra cAMP kit (Perkin-Elmer, Beverly, MA, USA). In brief, pure hiPSC-CM spheroids were dissociated by incubation with AccuMax^TM^ (PAA Laboratories, Cölbe, Germany) for 60 min at 37 °C and pipetting every 15 min. The cell pellet was resuspended in HBSS (Hank’s Balanced Salt Solution) containing 5 mM HEPES and 0.5 mM IBMX (pH 7.4). This cell suspension was then treated according to the manufacturer’s instructions (LANCE Ultra cAMP Kit, Perkin-Elmer). The cell suspension (5 μl containing approximately 1,000 cells) was added to 5 μl of agonist solution and allowed to incubate for 30 min at room temperature in an OptiPlate 384-well plate (Perkin-Elmer). The detection mix containing Eu-cAMP tracer and ULight-anti-cAMP was added and allowed to incubate for 1 h. The assay was read on a Flexstation 3 system (LANCE settings: 340 nm Ex/665 nm Em; Molecular Devices, Sunnyvale, CA, USA). The cAMP standards included with the kit and cell suspensions stimulated with HBSS (containing IBMX) only served as the standard curve and internal controls, respectively. After determining the IC_50_ of COA-Cl, we compared the cAMP levels after the application of several compounds (phosphate-buffered solution [PBS] as a control and Propranolol, Isoproterenol, SEW2871, Forskolin, or COA-Cl) using a Lance Ultra cAMP kit. The plate was then read using an Envision multi-label plate reader (Perkin-Elmer).

### The PDE activity assay of pure hiPSC-CM spheroids

Pure hiPSC-CM spheroids were pre-treated with COA-Cl or control (PBS) for 30 min. These spheroids were dissociated by incubation with AccuMax^TM^ for 60 min at 37 °C and pipetting every 15 min. After incubation, these cells in 300 µl lysis buffer were briefly centrifuged at 20,000×*g* for 10 min at 4 °C. The lysed cells were then incubated with the substrate cAMP at 37 °C for 10 min, and using a PDE activity assay kit (Abcam, Inc., Cambridge, MA), the PDE activity was measured by reading at OD 620 nm on an Envision multi-label plate reader.

### The histological analysis and immunohistochemistry of cardiac organoids

Samples were collected with collagen gels, fixed with 4% paraformaldehyde PBS (Wako, Osaka, Japan), and embedded in paraffin. Sections (4-6 µm) were mounted on microscope slides after dewaxing and rehydration. Antigen retrieval was performed in Dako pH 9 EDTA buffer (Dako, Kyoto, Japan) with a microwave. Sections were incubated with primary antibodies as follows: Mouse anti-CD31 antibody (1:20; REF MD0823, Dako), Rabbit anti-CD90 antibody (1:100; bs-10430R, Bioss Inc, Woburn, MA, USA), and Mouse anti-Troponin T antibody (1:100; ab8295; Abcam) were applied as the primary antibody for 16 h at 4 °C. We used a Dako EnVision Systems HRP kit (Dako) for the secondary antibody, and immunohistochemical tissue staining was performed with Nichirei-Histofine simple-stain MAX-PO (Nichirei, Tokyo, Japan). Hematoxylin was used for counterstaining.

### An immunofluorescence analysis of cardiac organoids

Samples were fixed in 4% paraformaldehyde phosphate-buffered solution for more than 16 h at 4 °C, blocked in Carbo-Free Blocking Solution (VECTOR LABOLATORIES INC., CA, USA) for 10 min, and permeabilized by 0.2% TritonX-100 (Wako, Japan) for 10 min at room temperature (20 °C). Primary and secondary antibody staining were performed as described below: Gout anti-troponin T antibody (1:100; C-18; Santa Cruz Biotechnology, CA, USA) was applied as the primary antibody for 16 h at 4 °C, while Gout Anti-Mouse IgG(H+L) and Alexa Fluor^®^ 488 (1:100; Thermo Fisher Scientific K.K., Tokyo, Japan) were applied as secondary antibodies for 60 min at 20 °C. DAPI

(Sigma–Aldrich Chemie, Steinheim, Germany) double-staining of the nucleic acids was finally performed for 10 min at 20 °C. The immunofluorescence signals were observed under a confocal laser scanning microscope (ZEISS LSM880 Confocal Microscope; Carl Zeiss, Oberkochen, Germany) using a ×40 oil immersion.

### Statistical analyses

All data are expressed as the mean ± standard deviation, and all statistical comparisons were determined by a one-way analysis of variance followed by Tukey’s test or a two-tailed Student’s *t*-test. All statistical analyses were performed using the Excel and JMP software programs (SAS Institute, Cary, NC).

## Results

### A histological analysis and immunohistochemistry of cardiac organoids

Phase contrast microscopy revealed the spheroid morphology of cardiac organoids (Fig. 2A). Spindle and polygonal cells were shown to be mixed in cardiac organoids (Fig. 2B). CD31-positive endothelial cells were scattered in cardiac organoids (Fig. 2C). Cardiac myocytes, shown as troponin T-positive cells, were located diffusely inside cardiac organoids (Fig. 2D). HDFBs were also distributed inside cardiac organoids (Fig. 2E). In addition, cardiac organoids were fixed with 4% paraformaldehyde were immunostained with troponin T to indicate the details of cardiac myocytes. The sample was observed with a scanning microscope (Fig. 2F). Fiber-like structures stained by troponin T indicated functional cardiac myocytes in cardiac organoids.

**Figure 2.**
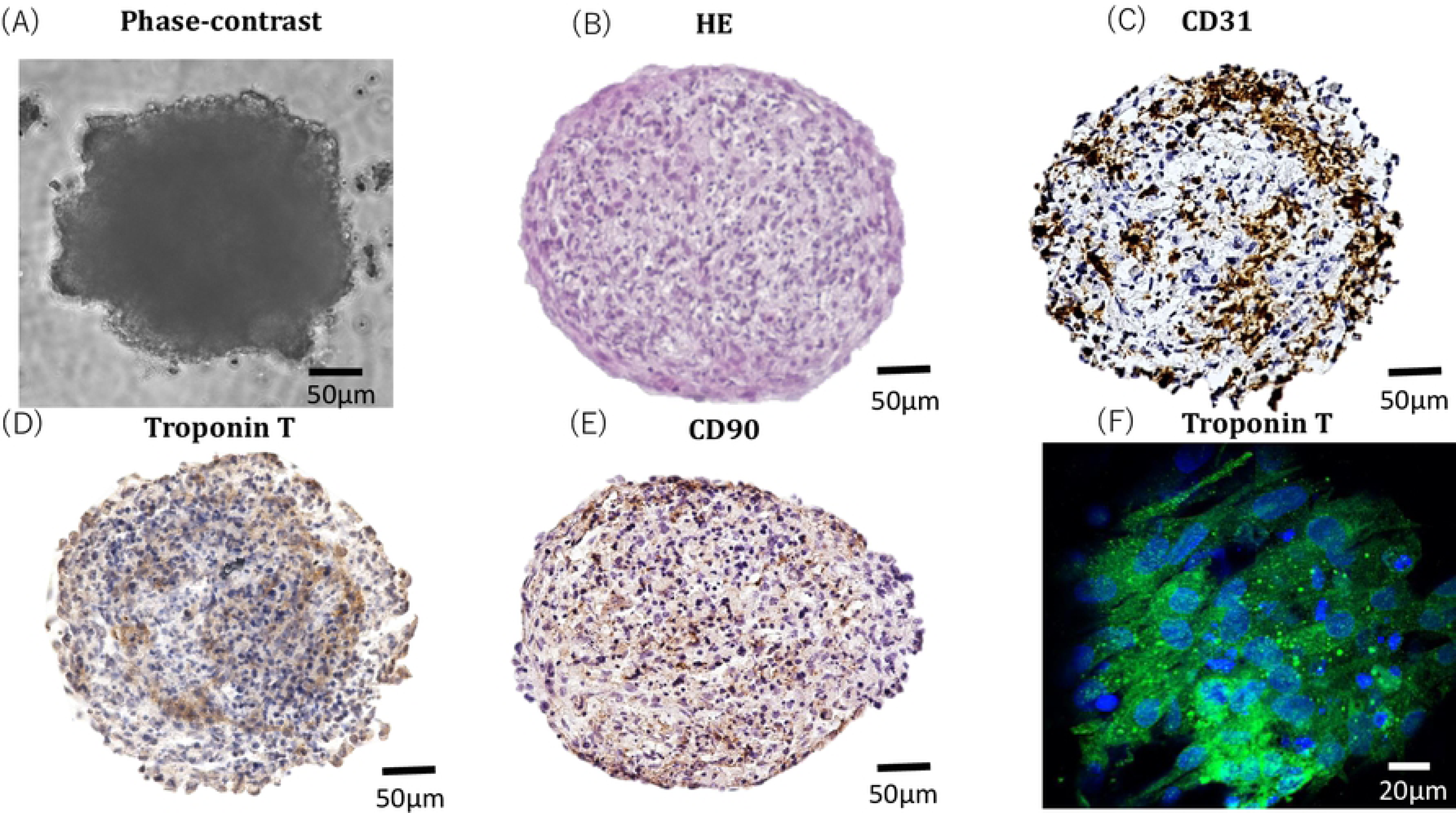
Phase contrast microscopy shows the spheroid morphology of cardiac organoids (A). Cardiac organoids were fixed with paraffin and subjected to HE staining (B). The representative images show immunostained cardiac organoids fixed in paraffin. Endothelial cells within the cardiac organoids are indicated by CD31 (C). Cardiac myocytes in cardiac organoids were stained with troponin T (D). The presence of functional cardiac myocytes in cardiac myocytes was indicated by troponin T staining. HDFBs in cardiac organoids were stained with CD90 (E). Cardiac organoids were fixed with 4% paraformaldehyde and subjected to immunofluorescence staining with troponin T. The sample was observed with a scanning microscope under a ×40 oil lens (F).

### Effects of COA-Cl on the beating and contraction of cardiac organoids

Research using disease models and drug screening systems with hiPSC-CMs is advancing rapidly. The tissue-engineered cardiac system with stem cells has become quite useful since the development of a more sensitive drug test model and an improved cell delivery strategy. The cardiac organoids in our study were composed of hiPSC-CMs, hiPSC-ECs, and HDFBs. First, they were immunostained with CD31, and tube formation was observed. Cardiac myocytes were also immunostained with Troponin T in order to observe the cardiac myocytes of the organoids in detail. HDFBs were also immunostained with CD90. Cardiac organoids fixed with 4% paraformaldehyde phosphate-buffered solution were also stained with troponin T antibody and observed with a confocal microscope.

Based on our findings, the cardiac organoids were determined to be a cardiac model with a vascular network. The effect of COA-Cl on the cardiac organoids was then investigated by observing this model before and after the addition of the drug. The movements of the cardiac organoids created and cultured for 10 days were observed. The beatings were recorded for 30 seconds. COA-Cl (1 mM) was then added, and the cardiac organoids were incubated for 30 min and recorded again. The fractional area change was determined from these recordings (Fig. 3A). The changes in the fractional area predominantly increased after the administration of COA-Cl. There was also a remarkable increase in the beating rate (10.7±0.7 beats/min, N=3), and COA-Cl (25.3±0.7 beats/min, N = 3) (Fig. 3B).

**Figure 3.**
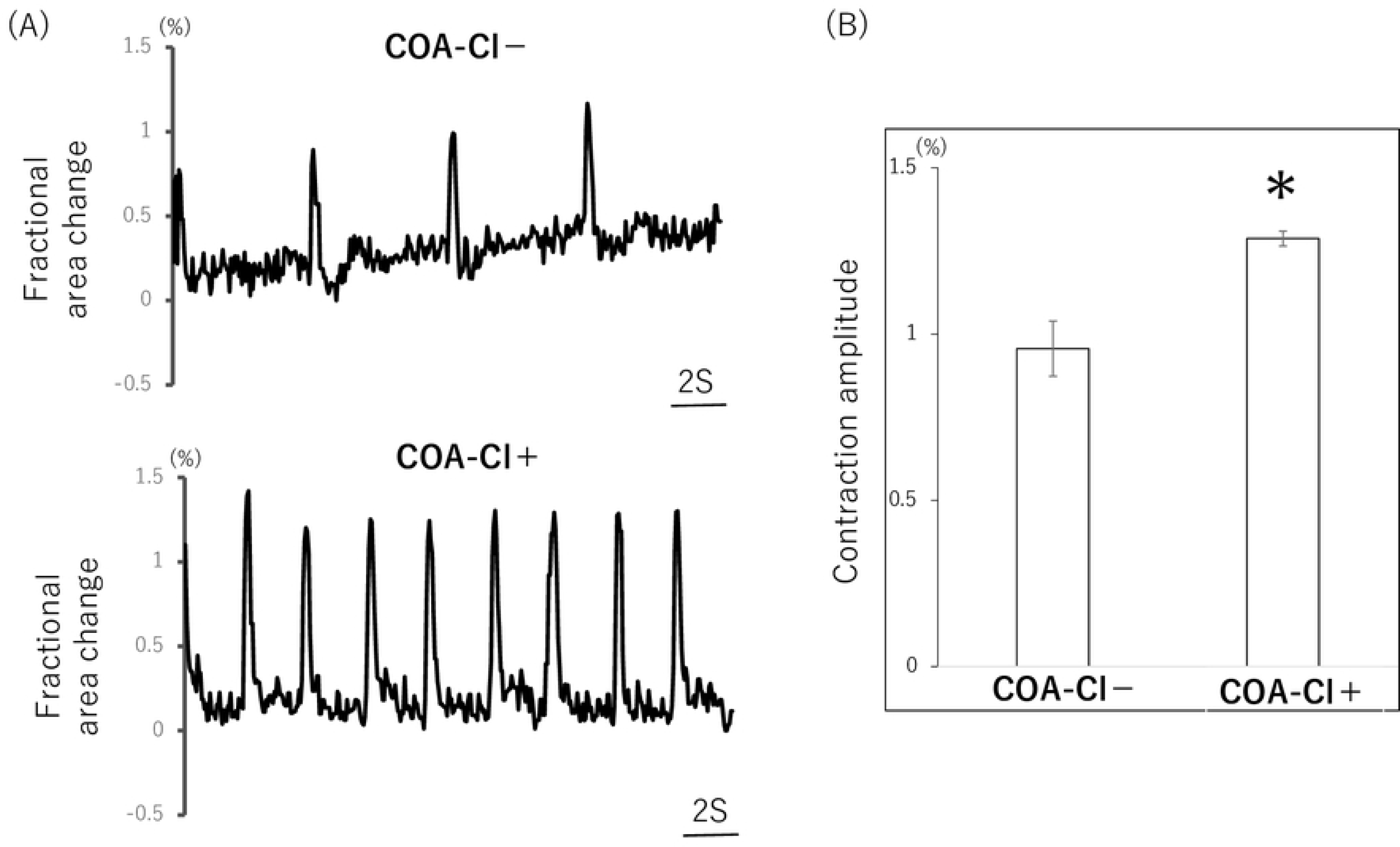
Cardiac organoids in bright-field images were observed on day 10. Representative contraction profiles of cardiac organoids were constructed based on a video analysis of the spheroid fractional area change (A). The analysis of the average (±SEM) contraction amplitude (i.e. fractional area change) showed significant differences on day 10 (n = 4 organoids). COA-Cl significantly increased the beating rate and the fractional area change in organoids (B). Asterisk indicates a statistically significant difference, p < 0.05.

### Effects of COA-Cl on the beating of pure hiPSC-CM spheroids

We tested the direct effects of COA-Cl on the beating rate of pure hiPSC-CM spheroids, which may be a valuable phenotypic parameter for drug discovery and development. A cardiac beating assay was performed to assess several positive and negative inotropic effects.

Positive chronotropic agents were Isoproterenol and Forskolin, which increase the amount of cAMP in cardiac myocytes and induce a positive inotropic effect. The negative inotropic agent was Propranolol, a nonselective β-blocker that inhibits the binding of norepinephrine (released from sympathetic nerve terminals) to the alpha-1 receptors. In addition, SEW2871 (S1P1 specific agonist) was used for the comparison with COA-Cl. The results from a representative experiment are shown in Fig. 4A. The beating rates of pure hiPSC-CM spheroids increased 30 min after the application of Isoproterenol and Forskolin. In contrast, these rates decreased after the application of Propranolol. SEW28171, an S1P1-specific agonist, did not increase the beating rate. The mean beating rates (±standard error of the mean) of the drug-free controls were as follows: vehicle (61.5±1.0 beats/min, N=4), Isoproterenol (91.0±0.6 beats/min, N=4), Propranolol (57.5±2.2 beats/min, N=4), Forskolin (99.5±0.5 beats/min, N=4), SEW2871 (62.5±1.0 beats/min, N=4), and COA-Cl (77.5±0.5 beats/min, N=4) (Fig. 4A). These results suggest that COA-Cl has the potential to increase the beating rate of pure hiPSC-CM spheroids, which is well known to indicate positive inotropic effects. However, the mechanism through which the beating is increased differs from that underlying the increase in beating that occurs with S1P1 activation.

**Figure 4.**
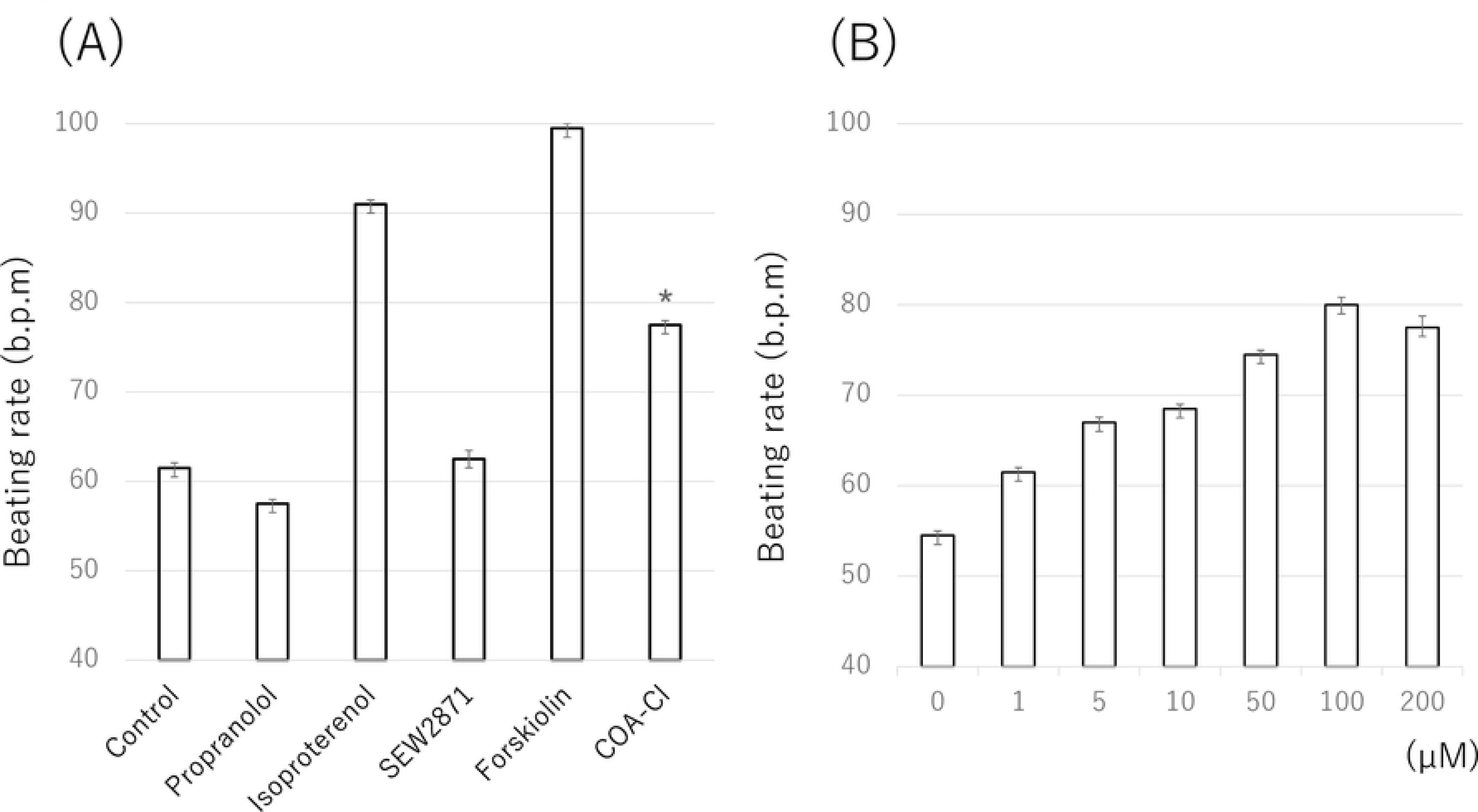
COA-Cl, ISO and FSK significantly stimulated the beating rates. PRP suppressed the beating rate. No marked changes were observed with SEW2871. The data were obtained from four individual experiments, each of which was performed four times. The average values (±SEM) normalized with reference to the basal condition in the presence of PBS (A). COA-Cl stimulated the beating rates in a concentration-dependent manner. Seven different concentrations of COA-Cl were prepared: 0 M (baseline), 1 μM, 5 μM, 10 μM, 50 μM, 100 μM, and 200 μM. These spheroids were exposed to these compounds for 30 min and recorded for 30 s using a BZX-710. The data were obtained from four individual experiments, each of which was performed four times. The average values (±SEM) were normalized with reference to the basal condition in the presence of PBS (B).

We also observed the concentration-dependent changes in the beating rate. To evaluate the effective concentration of COA-Cl for pure hiPSC-CM spheroids, the spheroids were treated with increasing concentrations of COA-Cl (0-200 µM). Results from a representative experiment are shown in Fig. 4B. A statistical analysis of the beating rates showed a significant increase at all. COA-Cl concentrations. To detect the onset of the beating change with the administration of COA-Cl, calcium staining was performed in order to measure the rate of signal waves through beating spheroids. Pure hiPSC-CM spheroids were treated with fluo-4, a calcium-binding fluorescent dye, and visualized by fluorescence microscopy. The trace of cell-beating disruption after the addition of COA-Cl is shown in Fig. 5. After the administration of COA-Cl (1 mM), the beating rate was 1.5-fold faster than before the administration. These results show that COA-Cl is strongly related to some types of receptor and ion channel phosphorylation because the effects occurred immediately after its administration.

**Figure 5.**
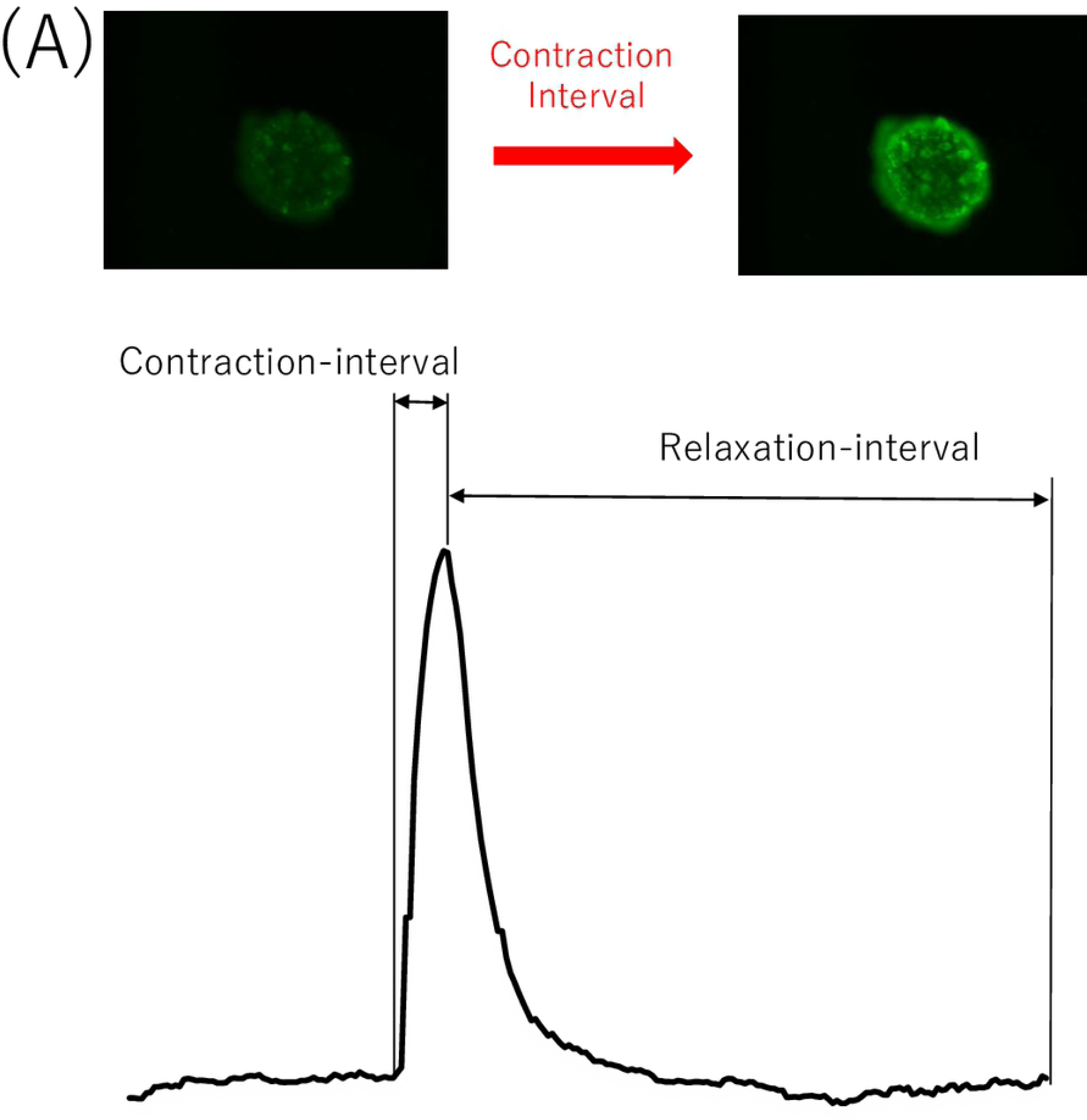
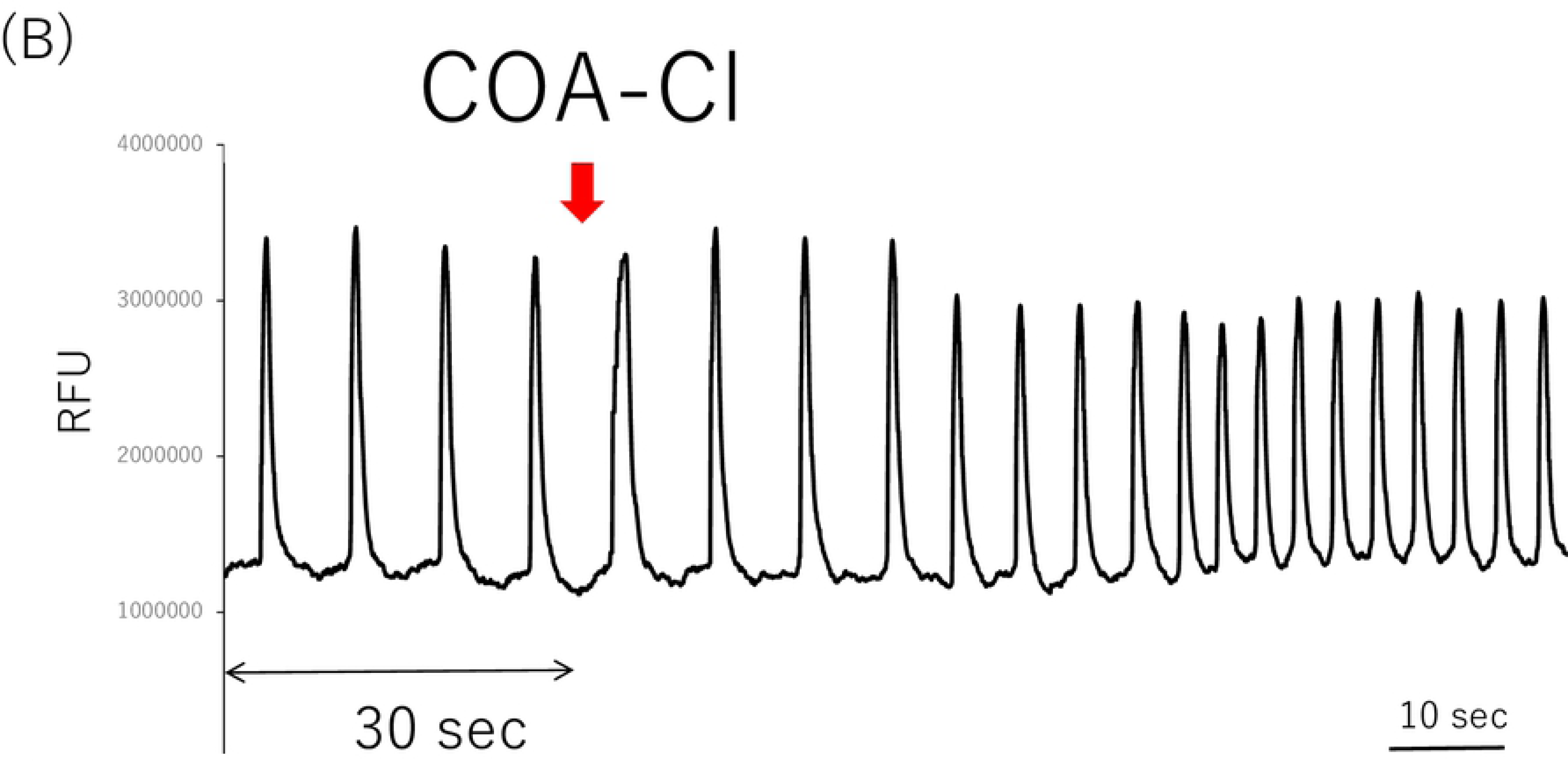
The contraction of pure hiPSC-CM spheroids was visualized using Fluo-4 (a calcium indicator). Pure hiPSC-CM spheroids were then loaded with Fluo-4, which fluoresces upon calcium binding. Representative still images of resting and contracting pure hiPSC-CM spheroids, taken from a time-lapse video capturing the calcium influx during contraction. Rate-interval graph demonstrating spheroid contraction as measured by recording the changes in the Fluo-4 fluorescence intensity. The contraction interval was determined as the time between low Fluo-4 fluorescence (relaxed cells) and high Fluo-4 fluorescence intensity (contracted spheroids) (A). COA-Cl increased the beating rate of pure hiPSC-CM spheroids in the early phase. The medium was loading buffer. Pure hiPSC-CM spheroids were assessed to determine their ability to contract using a Fluo-4 calcium binding assay. The rate interval graph shows fluctuations in the Fluo-4 fluorescence intensity, indicative of pure hiPSC-CMs spheroid contraction, under basal conditions for 30 s and after sequential treatment with COA-Cl. COA-Cl affected the rate of contraction at an early stage. The contraction rate of differentiated pure hiPSC-CM spheroids was determined as the number of contractions (contraction interval + relaxation interval) per minute. The data were from nine individual experiments, each of which was performed three times (B).

Overall, these data demonstrate that pure hiPSC-CM spheroids mimic the cellular and molecular composition of human heart tissue. Thus, the beating rate of pure hiPSC-CM spheroids may be a valuable phenotypic parameter for use in drug discovery and development. COA-Cl was found to have positive inotropic effects with pure hiPSC-CM spheroids.

### Effects of COA-Cl on the cAMP lefels of pure hiPSC-CM spheroids

The baseline cAMP accumulation of hiPSC-CMs was evaluated with increasing concentrations of Forskolin (100 mM to 1 nM, half-log increments) and time-resolved fluorescence resonance energy transfer (TR-FRET). cAMP standards were used to construct a TR-FRET-concentration curve and calculate the amount of cAMP (Fig. 6A). The baseline cAMP concentrations, as determined by the Forskolin method, showed a dynamic range of 1 nM to 100 µM (Fig. 6B). Subsequently, the effects of COA-Cl were evaluated. COA-Cl (1 µM to 100 mM; half-log increments) caused a concentration-dependent decrease in TR-FRET and thus an increase in cAMP. In contrast, SEW2871 (1 pM to 1 µM; half-log increments) caused no concentration-dependent change in TR-FRET. These data indicate that COA-Cl increases the cAMP concentration in pure hiPSC-CM spheroids via a different mechanism from SEW2871, and we determined that the IC_50_ of COA-Cl was 1 mM. We then compared the results with vehicle (PBS), Isoproterenol IC_50_, Forskolin, Propranolol, and SEW2871. Forskolin, ISO, and COA-Cl showed additive effects in the total cAMP produced. In contrast, Propranolol and SEW2871 had no remarkable effect on the total cAMP production. These data were very similar to those obtained in the beating analysis. COA-Cl has also been reported to bind to the A1R; however, its action was not clear. We confirmed that COA-Cl increases the cAMP levels of hiPSC-CMs through the A1R. These cells were incubated with Suramin (100 μM) for 1 h followed by incubation with several compounds and measurement of the cAMP levels. These data indicated that the inhibition of the A1R by Suramin was not involved in the increase in the cAMP concentrations of hiPSC-CMs (Fig. 6E, 6F).

**Figure 6.**
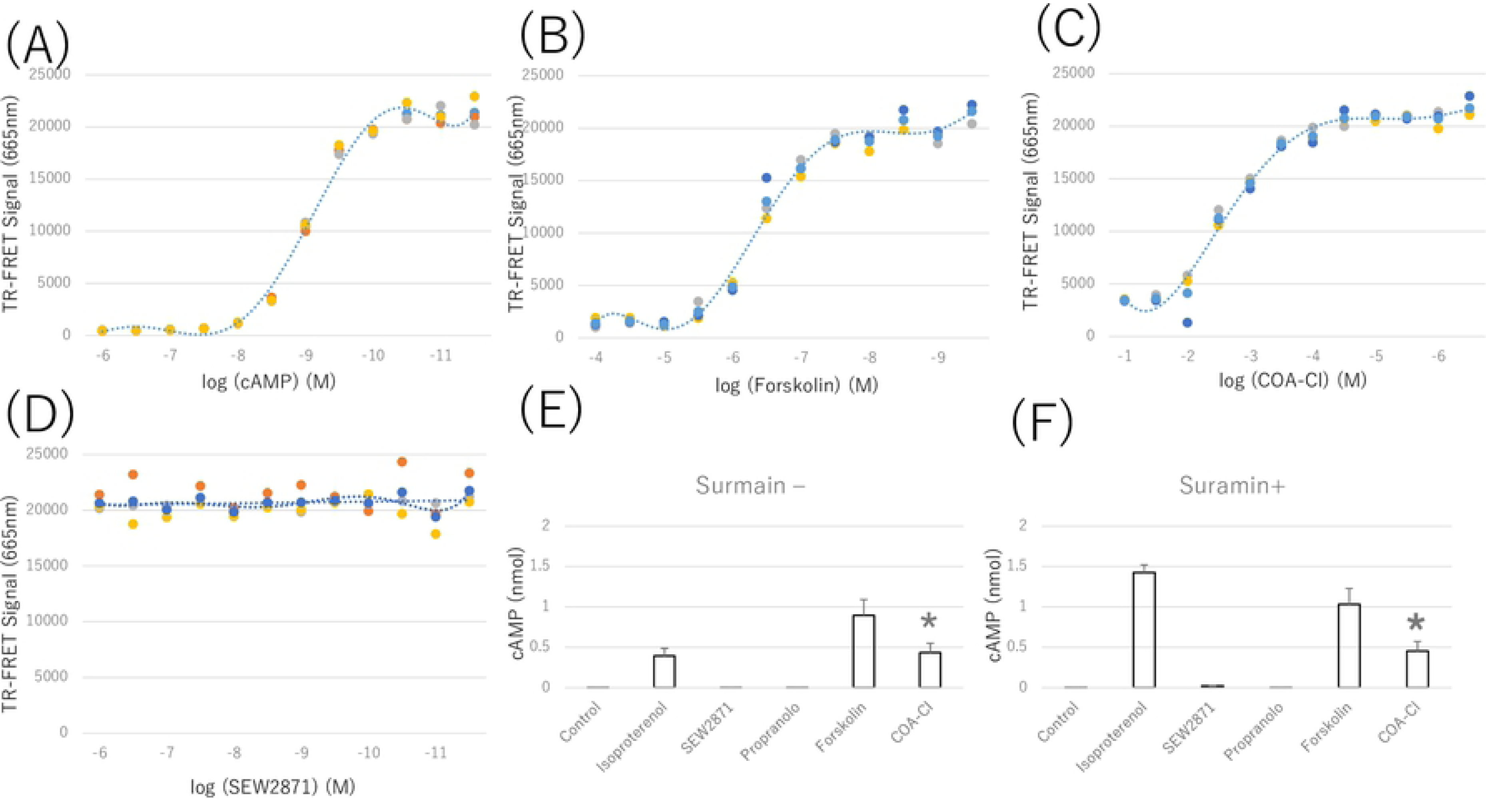
cAMP standards (10 pM-1 μM; half-log increments) were used to create a concentration calibration based on the decreasing time-resolved fluorescent resonance energy transfer (TR-FRET) in hiPSC-CMs (A). Forskolin (1nM-100 μM; half-log increments) caused a concentration-dependent decrease in TR-FRET and thus an increase in cAMP. The dose response curve was drawn using the mean values (B). COA-Cl (100 μM-10M; half-log increments) caused a concentration-dependent decrease in TR-FRET and thus an increase in cAMP. The dose response curve was drawn using the mean values (C). In hiPSC-CMs, COA-Cl (1 mM), ISO (10 nM), and FSK (10 µM) substantially increased the cAMP level compared to the basal cAMP level in untreated controls. Treatment with SEW2871 (20 nM) and PRP (10 nM) did not significantly alter the cAMP levels compared to control. *P values of < 0.05 were considered to represent a statistically significant difference compared to control (n = 5–8) (D). HiPSC-CMs treated with Suramin (100 µM), COA-Cl (1 mM), ISO (10 nM), and FSK (10 µM) showed significantly increased cAMP levels compared to the basal cAMP level in untreated controls. Treatment with SEW2871 (20 nM) and PRP (10 nM) did not significantly alter the cAMP level compared to control. *P values of < 0.05 were considered to represent a statistically significant difference compared to control (n = 5–8) (E). *P < 0.05 significant compared to control. These results showed that COA-Cl increased the cAMP levels of hiPSC-CMs similarly to Isoproterenol and Forskolin, but not through A1R or S1P1R activation (n = 5–8).

In conclusion, the mechanism through which COA-Cl increases the cAMP levels does not involve S1P1R or A1R, which are known to be COA-Cl-binding receptors. COA-Cl increased the cAMP levels of hiPSC-CMs similarly to Isoproterenol and Forskolin.

### Effects of COA-Cl on PDE in pure hiPSC-CM spheroids

Previous studies have shown that xanthine derivatives act as nonspecific inhibitors of PDE. The action of xanthine derivatives is mainly due to the increase in the intracellular cAMP concentration as a second messenger by the inhibition of PDE. Theophylline is a powerful bronchodilator that is used to treat respiratory diseases, such as COPD (chronic obstructive pulmonary disease) [16]. It is also known to have a positive inotropic effect on the heart [17]. COA-Cl is a nucleic acid analog that is structurally similar to a xanthine derivative (Fig. 7A).

**Figure 7.**
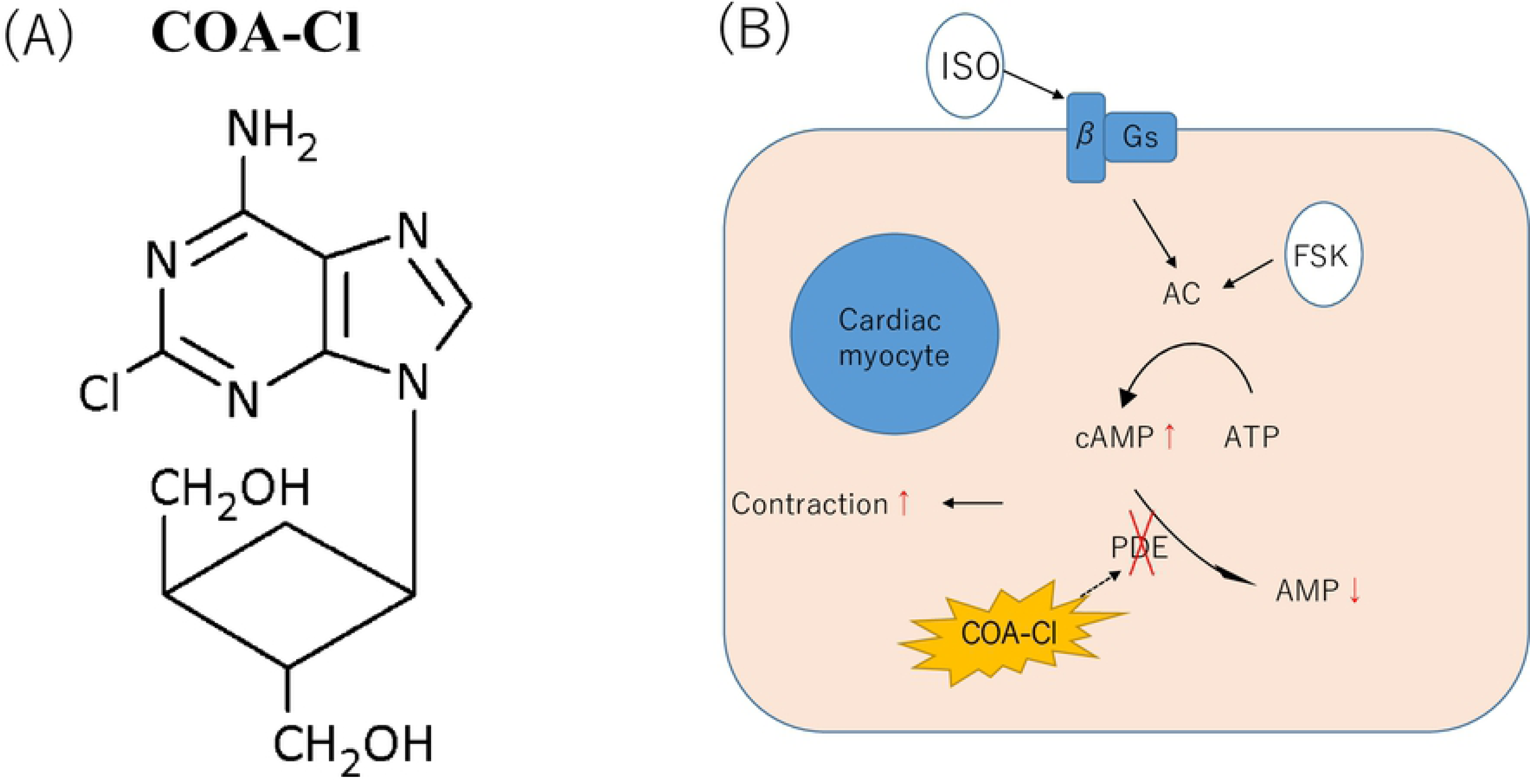
A schematic illustration showing the structure of COA-Cl [3] (A). A schematic model of cardiac excitation-contraction coupling through PKA. Possible sites for the regulation of excitation-contraction coupling by the AC/PKA cascade are indicated (B).

We hypothesized that COA-Cl was involved in the elevation of cAMP in cardiac myocytes through its action as a PDE inhibitor. To explain the cause of cAMP elevation, we investigated the activity of the PDE using hydrolyzing cAMP time-course and found that the administration of COA-Cl reduced the PDE activation in comparison to control in hiPSC-CMs (Fig. 8). This result suggests that COA-Cl may promote the cAMP elevation of hiPSC-CMs and is involved in the increasing pulsation of pure hiPSC-CM spheroids.

**Figure 8.**
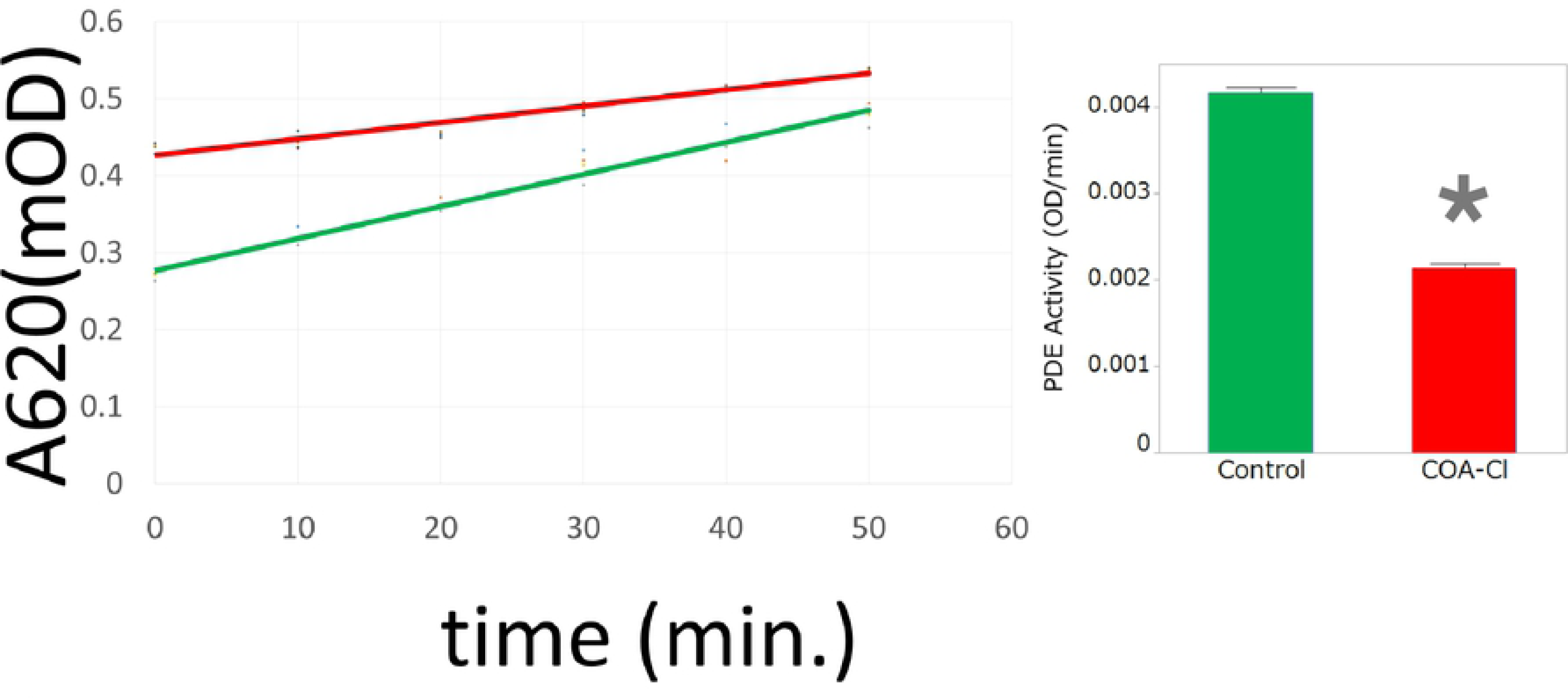
COA-C significantly induced PDE activity in hiPSC-CMs. PDE enzyme (20 mU/well) was incubated at 30 °C for the indicated times with cAMP (200 μM) and 5’-nucleotidase (50 kU/well) with or without COA-Cl (1 mM). Reactions were terminated by the addition of 100 µL of Green Assay Reagent and OD 620 nm read 30 min later. A standard curve of cAMP can be used to convert the OD 620 nm data to nmol of 5’-AMP. Each data point represents the mean and SEM of three independent experiments performed in triplicate.

**Figure 9.**
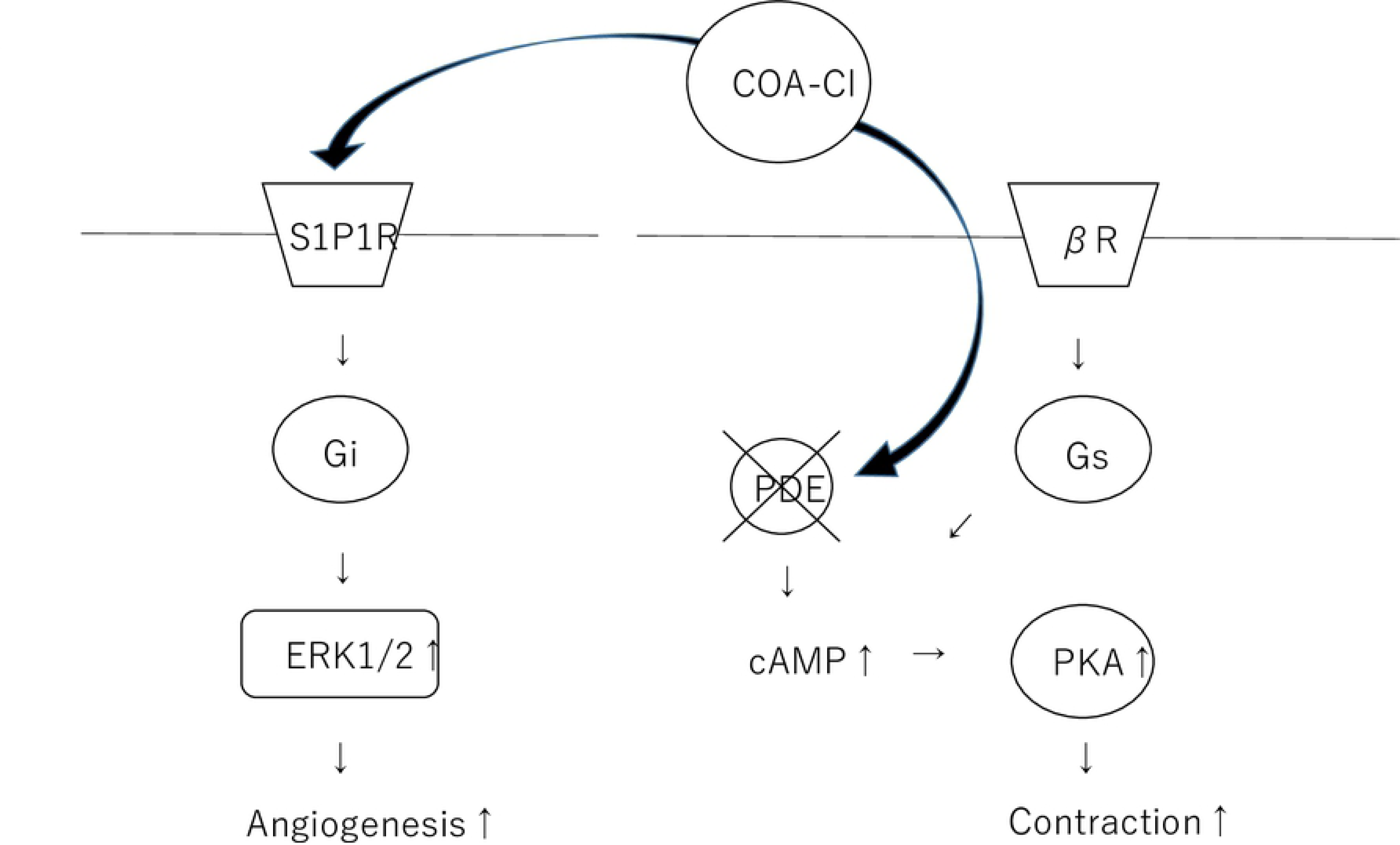
A schematic illustration showing the cascade of both angiogenesis and the positive inotropic effects.

## Discussion

In this study, we showed that COA-Cl suppresses PDE and increases the contraction of cardiac organoids, independent of S1P1R and A1R. COA-Cl can produce signaling molecules and tube formation in cultured human vascular endothelial cells through S1P1R-mediated extracellular stimulation [3]. COA-Cl has been shown to exert a neuroprotective effect against intracerebral hemorrhaging [6]. It has also been reported that COA-Cl may reduce oxidative stress, which may be one of the mechanisms underlying its neuroprotective effect [5]. In addition, in the perfused heart, pretreatment with S1P has been shown to significantly restore the heart function after ischemia. Interestingly, the protective effect of ischemic preconditioning on ischemia/reperfusion injury was promoted in the hearts of rats treated with S1P1R and S1P3R antagonists [18][19].

COA-Cl is a small molecule that is slightly similar to adenosine. It was hypothesized that it may bind to the G protein-coupled receptor (GPCR) rather than receptor tyrosine kinase (RTK). COA-Cl was shown, through an exhaustive investigation, to act as an agonist of S1P1R and to bind to adenosine A1R [3]. However, the effects of A1R are still unknown. COA-Cl is considered to be a partial stimulator of S1P1R. S1P1R couples with the Gi family of guanine nucleotide-binding regulatory proteins (G-proteins) to activate multiple intracellular signaling pathways, including the extracellular signal-regulated kinase 1, 2 (ERK 1/2) pathways [3]. It has been shown that S1P1R activates sprouting angiogenesis and maintains blood vessel stability in endothelial cells, while S1P1R antagonism inhibits tumor vascularization [20] [21]. However, the mechanisms underlying these potent effects of COA-Cl have remained unclear.

To facilitate drug discovery, we created cardiac organoids and then predicted the effects of COA-Cl on the heart and analyzed these functions using the cardiac organoids. We discovered that COA-Cl enhances the pulsatile and contractile power and that increases the cAMP level via the partial inhibition of PDE. Clinically, PDE inhibitors are positive inotropic drugs that are useful for treating heart failure. These contractile actions ultimately increase the Ca^2+^ release of the sarcoplasmic reticulum (PDE) by activating cAMP-PKA. It has also been reported that the cAMP and cGMP-PDE activities after ischemic preconditioning are higher than in non-preconditioned hearts and hearts with elevated levels of cGMP [22]. PDE inhibitors are expected to have clinical application as drugs that improve the cardiac function under severe conditions, such as ischemia. Numerous studies have shown that positive inotropic drugs exacerbate the prognosis. Whether or not COA-Cl impairs the prognosis as well as the patient’s quality of life is unclear; however, it may improve the cardiac output in order to escape the vicious cardiac cycle and condition characteristic of the decompensated phase of heart failure.

Spheroid formation is a well-known phenomenon that occurs based on the nature of cell aggregation. It was reported that isolated beating ventricular heart cells grown in tissue culture showed a tendency to beat in three dimensions from single-layer cell sheets [23]. Compared with monolayer cultured cells, adhesion cells or hepatocyte spheroids can be observed in conditions that more closely resemble those of living bodies, which is useful for drug screening of anticancer drugs and evaluating the liver function [24][25].

There has been a report on the application of spheroids to promote the secretion of dopamine in a Parkinson’s disease model [26]. A cell matrix provides a very important framework that allows cells to function and release bioactive substances (nutritional ingredients) in order to maintain the structure of tissue compounds [27]. When creating cardiac organoids, it is difficult to make them without certain materials, such as hydrogel [28]. We made cardiac organoids without any materials. We also successfully fabricated a 3D tubular structure made of spheroids with a scaffold-free bio 3D printer to make small-caliber vascular prostheses [13]. Studies of scaffold-free bio 3D printers have been conducted in various fields [29] [30]. Takahashi and Yamanaka reported a method for reprogramming fully differentiated fibroblasts derived from the tissues of adult or fetal mice to make cells similar to ES cells [31]. The availability of human cardiac myocytes with differentiated pluripotent stem cells offers a new opportunity to construct *in vitro* models of heart disease [32], conduct drug screening for new drugs [33], and apply cardiac therapy to specific patients [34].

The application of hiPSC-derived cells to tissue engineering has also been a focus of research. Since the presence of a cell matrix strongly influences the beating of the heart muscle, we decided to create an organoid that mimics the heart with fibroblasts and endothelial cells. These organoids therefore reflect the heart *in vitro*. However, there are few reports on the partial shortening of the 3D cardiac structure without scaffolds. Partial shortening was observed in a study of Mukae et al., wherein the authors monitored 2 points (using a software program) with a 2D cardiac patch, and approximately 2%–3% area changes was observed [35]. Richards et al. compared the adult-stage heart with the developing-stage heart by making cardiac organoids and measuring their fractional change [36]. We investigated the contractile action in a human cardiac organoid model with and without COA-Cl by observing the fractional area change and the beating rate of cardiac organoids. Using these techniques, we prepared mixed spheroids derived from hiPSCs and evaluated the drug effect on the cardiac organism based on the beating rate and area change.

In our study, we examined the effect of the novel nucleic acid analogue COA-Cl on the heart. In order to analyze cardiotonic drugs *in vitro*, cardiac organoids composed of hiPSC-derived cells were constructed. COA-Cl resulted in pulsatility and contraction changes in the human heart model. Similarly to other cardiotonic drugs, COA-Cl increased the intracellular cAMP level. It was confirmed that its PDE inhibiting effect was one factor that increased the cAMP level.

COA-Cl is a xeno-free compound and may therefore be safely applied in regenerative medicine. The growth of cells *in vitro* usually involves a basal medium supplemented with FBS, which is associated with a risk of disease transmission. Furthermore, growth factors such as VEGF and FGF are fragile and may be subject to considerable lot-to-lot variability [37]. Thus, in tissue engineering with transplantation in mind, the culture of tissues under serum-free condition with substances that are not antigenic is required. As COA-Cl is a stable nucleic analogue, it can be applied to tissue engineering for regenerative medicine.

## Conclusion

COA-Cl was shown to act as a partial PDE inhibitor of cardiac organoids. Cardiac organoids are useful for drug discovery, facilitating assay performance. The results of the present study suggest their possible application in cardiac transplantation for regenerative medicine. COA-Cl can be used as a cardiac inotropic agent, as a partial PDE inhibitor, and to promote angiogenesis in patients with ischemic heart disease and heart failure.

## Acknowledgements

This research was supported by JSPS KAKENHI Grant Number 16H02674, 17H04292, 17K10732. None of the authors has a financial relationship with a commercial entity that has an interest in the subject of the present manuscript or other conflicts of interest to disclose. The authors are grateful to H. Nakao, I. Nishioka, H. Tsugitomi, H. Manako, and S. Haraguchi for their excellent technical assistance. The authors would also like to take this opportunity to thank. K. Oba, S. Matsuhashi, J. Xia, K. Mori, and Y. Tokuyama for their advice and for teaching the skills required to perform the experiments.

